# Allos: an integrated Python toolkit for isoform-level single-cell and spatial in-situ transcriptomics

**DOI:** 10.64898/2026.03.24.713944

**Authors:** Eamon McAndrew, Anna Diamant, Georges Vassaux, Pascal Barbry, Kevin Lebrigand

## Abstract

Single-cell RNA sequencing and spatial transcriptomics have transformed our understanding of the transcriptional landscape by enabling high-resolution profiling of gene expression. Yet most experimental pipelines and their associated analysis frameworks collapse transcript diversity into gene-level counts, obscuring alternative splicing and isoform usage. The increasing ability of long-read sequencing to recover full-length transcripts from single cells and spatially barcoded tissues has created a pressing need for computational frameworks to support the storage, analysis, and visualisation of isoform-resolved data. Existing tools for isoform and splicing analysis either specialise in bulk, single-cell, or spatial RNA-seq assays in isolation and remain fragmented across languages and data models, limiting interoperability and hindering widespread adoption. We present Allos, a Python framework for isoform-level single-cell and spatial transcriptomics analysis. Built on the AnnData data model, Allos natively represents transcript-level quantification and integrates directly with GTF/GFF and FASTA annotations. Allos enables differential isoform usage screening, multi-panel visualisation, structural transcript interpretation, and protein-level analysis across bulk, single-cell, and spatial assays from both long- and short-read sequencing. Its modular design and scverse compatibility allow isoform-resolved analyses to run alongside established gene-level workflows, linking transcript-level screening with structure-aware visualisation and downstream interpretation.

Allos is open-source and available at https://github.com/cobioda/allos, with comprehensive documentation and tutorials provided online.

## Background

### Isoform diversity in single-cell and spatial transcriptomics

Single-cell RNA-seq (scRNA-seq) and spatial transcriptomics (ST) have reshaped transcriptomics by resolving cellular heterogeneity and tissue architecture through high-resolution gene expression profiling of individual cells and of physical space within tissue sections (1–4). The vast majority of current scRNA-seq and ST approaches aggregate transcripts from each gene into a single count, thereby collapsing isoform diversity into a single feature per gene and providing an incomplete representation of the transcriptional landscape (5–7). This simplification has undoubtedly proven useful, but as protocols and sequencing increasingly enable the generation of high-resolution, full-length isoform data(8–14), computational frameworks to analyse these granular data types, tailored to the specific needs of single-cell and spatial isoform-resolution data, are needed (15–17).

Alternative splicing (AS) shapes gene regulation by producing distinct mRNA isoforms, tuning gene activity across tissues, cell types, developmental stages, and cellular states (18–20). Which isoforms are expressed depends on a combination of factors (disease state, tissue of origin, genetic background, and others), making splicing one of the more context-sensitive layers of gene regulation(21–26). Aberrant splicing underlies an estimated 15–30% of inherited human disorders (27–29). It is a well-established feature of cancer, where altered splicing programmes produce isoforms that drive proliferation, survival, invasion, and therapy resistance (30).

These isoform-level differences can have profound functional consequences independent of changes in gene expression, manifesting across a hierarchy of molecular levels. At the RNA level, sequence variation shapes transcript stability, subcellular localisation, and regulatory interactions with RNA-binding proteins and miRNAs (31–35). At the protein level, alterations in coding sequence can reshape domain architecture, eliminate or introduce post-translational modification sites, redirect subcellular targeting, or disrupt interaction interfaces. These changes may rewire entire signalling pathways, shift a protein between activating and inhibitory roles, or confer entirely new functions, all without any change in gene dosage (36–39). AS is but one method contributing to proteome diversity, but along with post-translational modifications and genetic variation, estimates suggest that the ∼20,300 protein-coding genes in the human genome give rise to more than one million distinct protein products upon translation (40). Full-length transcript sequences enabled by long-read sequencing that recapitulate transcript start sites, end sites, and exon–exon junction architecture are a critical resource for understanding the expansive proteoform landscape of individual cells (41,42).

### Long-read sequencing and the isoform-resolution data landscape

Existing single-cell and spatial analysis frameworks, such as Seurat and Scanpy, although widely used, are not explicitly designed to analyse transcript-level variation (43,44). Such frameworks were designed to handle data generated by standard short-read 3′ or 5′ tag-based protocols, such as those proposed by 10x Genomics. These technologies capture and sequence only the 3′ or 5′ ends of transcripts, often following fragmentation or non-full-length library generation, which introduces strong positional coverage biases that obscure internal exon structure and make it impossible to resolve full-length isoforms(5,7,45). Long-read (LR) sequencing technologies, such as Oxford Nanopore Technologies (ONT) and Pacific Biosciences (PacBio), have the potential to resolve full-length transcripts and novel isoforms in single-cell and in situ capture assays. Continuing improvements in long-read accuracy (46), coupled with advances in single-cell (11,47–49) and spatial (50) experimental protocols and dedicated computational pipelines, have made isoform-resolved quantification increasingly practical. Most workflows combine barcode and UMI recovery with transcript model construction and isoform-level quantification, producing cell- or spot-by-isoform count matrices. Several pipelines now support this, including SiCeLoRe, FLAMES, Isosceles, SCOTCH, bambu/bambu-clump, and lr-kallisto (11,12,51–54).

In parallel, both ONT and PacBio distribute and maintain platform-specific analysis workflows. For ONT, the EPI2ME Labs wf-single-cell workflow is an ONT-supported pipeline for processing Nanopore single-cell and spatial libraries (including Visium and Visium HD), covering barcode/UMI extraction and initial downstream analysis. For PacBio, the Iso-Seq tool suite is integrated into the SMRT Link/SMRT Analysis software for processing long-read transcriptome datasets (55,56).

A broad ecosystem of existing software supports the analysis of alternative splicing, but tools are often specialised for particular assay types and analysis units. For bulk RNA-seq, differential transcript/isoform usage can be tested with frameworks such as IsoformSwitchAnalyzeR and BANDITS, while event-based approaches such as rMATS and SUPPA2 quantify changes in specific splicing events (57,58). A smaller set of methods targets long-read transcript models and isoform visualisation (e.g., Swan, ScisorWiz), and several approaches address isoform usage estimation in single-cell contexts (e.g., BRIE) (59–61). In practice, approaches designed for bulk differential isoform usage models (including DEXSeq, DRIMSeq and satuRn) are also commonly applied to single-cell studies after aggregation (pseudobulking/metacells) (62–64). Across these categories, tooling typically operates on different primary representations (event counts, transcript-level abundances, transcript models, or aggregated matrices). It integrates with different downstream workflows, which can make it non-trivial to combine end-to-end isoform discovery, quantification, testing and visualisation within a single analysis environment, particularly for Python/AnnData-based pipelines. Recent benchmarking studies highlight key strengths and persistent limitations of long-read isoform analysis workflows, spanning transcript identification, quantification, and reproducible method comparison across platforms and protocols. Recent benchmarking studies reinforce this picture, demonstrating that method performance varies considerably by protocol, platform, and analysis task, and pointing to a clear need for reproducible, interoperable end-to-end workflows for long-read isoform analysis across both bulk and single-cell/spatial settings. (65,66).

### Towards a Python-native framework for isoform-level analysis

The success of gene-level single-cell analysis has been driven largely by the emergence of integrated analysis frameworks, which define shared data representations and support rich ecosystems of interoperable tools(43,44). These frameworks have become the default entry point for users analysing single-cell data, enabling rapid exploratory analysis while supporting extensibility and method development at scale. Although numerous studies have demonstrated the power of isoform-resolved single-cell analysis (67–70), often through adaptations of existing tools or bespoke, task-specific pipelines, these approaches remain fragmented. The development of a unified, user-friendly, and extensible framework for isoform-level analysis, therefore, represents an important and timely research direction.

To address these limitations, we present Allos, a Python package that provides an end-to-end solution for isoform-level analysis in single-cell and spatial in-situ transcriptomics. Allos is agnostic to the upstream quantification pipeline and supports transcript × cell or spot count matrices from various platforms. While Allos supports isoform-resolution data from multiple platforms, its design was motivated by and optimised for long-read single-cell sequencing data, where full-length transcript information is available. Extending the AnnData framework, Allos implements differential isoform usage testing. Allos emphasises screening based on per cent spliced in (PSI), a widely used metric quantifying the relative abundance of each transcript as a proportion of total gene expression, coupled with specialised visualisation of isoform architecture. Allos facilitates scalable, interpretable analysis of both known and novel transcriptomes, offering an integrated environment for exploring isoform diversity at bulk, single-cell, and spatial resolutions.

## Results

### Allos provides generalised quality control for isoform-resolution datasets

Any isoform-resolution analysis should begin with data quality assessment, given the limitations that can arise from sample quality, library preparation, and the lack of standardised long-read protocols. To demonstrate Allos, we use two datasets throughout this study. The first profiles the hippocampus, cortex, and ventricular zone of an E18 C57BL/6 mouse using ScNaUmi-seq, which combines 10x Genomics Chromium capture with long-read sequencing and Illumina-guided barcode assignment, covering over 1,000 cells across major neuronal and progenitor populations (11). The second comprises two adult (P56) mouse coronal brain sections (CBS1 and CBS2), generated using the SiT framework, pairing 10x Genomics Visium with long-read sequencing to quantify isoforms at each capture spot while retaining tissue architecture (17).

Allos provides a set of QC plots to assess isoform-resolution data rapidly before downstream analysis. Applied to the E18 dataset, these cover UMIs per cell (**Fig.2a**), cross-platform concordance with matched Illumina short reads (**Fig.2b**), isoform complexity per gene (**Fig.2c**), and transcript length distributions weighted by abundance (**Fig.2d**). Cross-platform concordance and isoform complexity plots can be facetted by cell type, sample, condition, or batch. The concordance module also supports gene- and isoform-level matrix comparisons from pipelines that output both, including ONT and PacBio workflows, as well as gene-detection differences between spatial platforms such as Visium and Xenium. On the CBS spatial dataset, gene body coverage from BAM files revealed the 3′ bias expected from 3′ capture protocols (**Fig.2e**), with consistent profiles across CBS1 and CBS2 (3′/5′ ratios of 2.12 and 1.91), confirming reproducible library preparation. Together, these diagnostics serve as a practical quality checkpoint for any long-read single-cell or spatial dataset.

**Figure 1.**
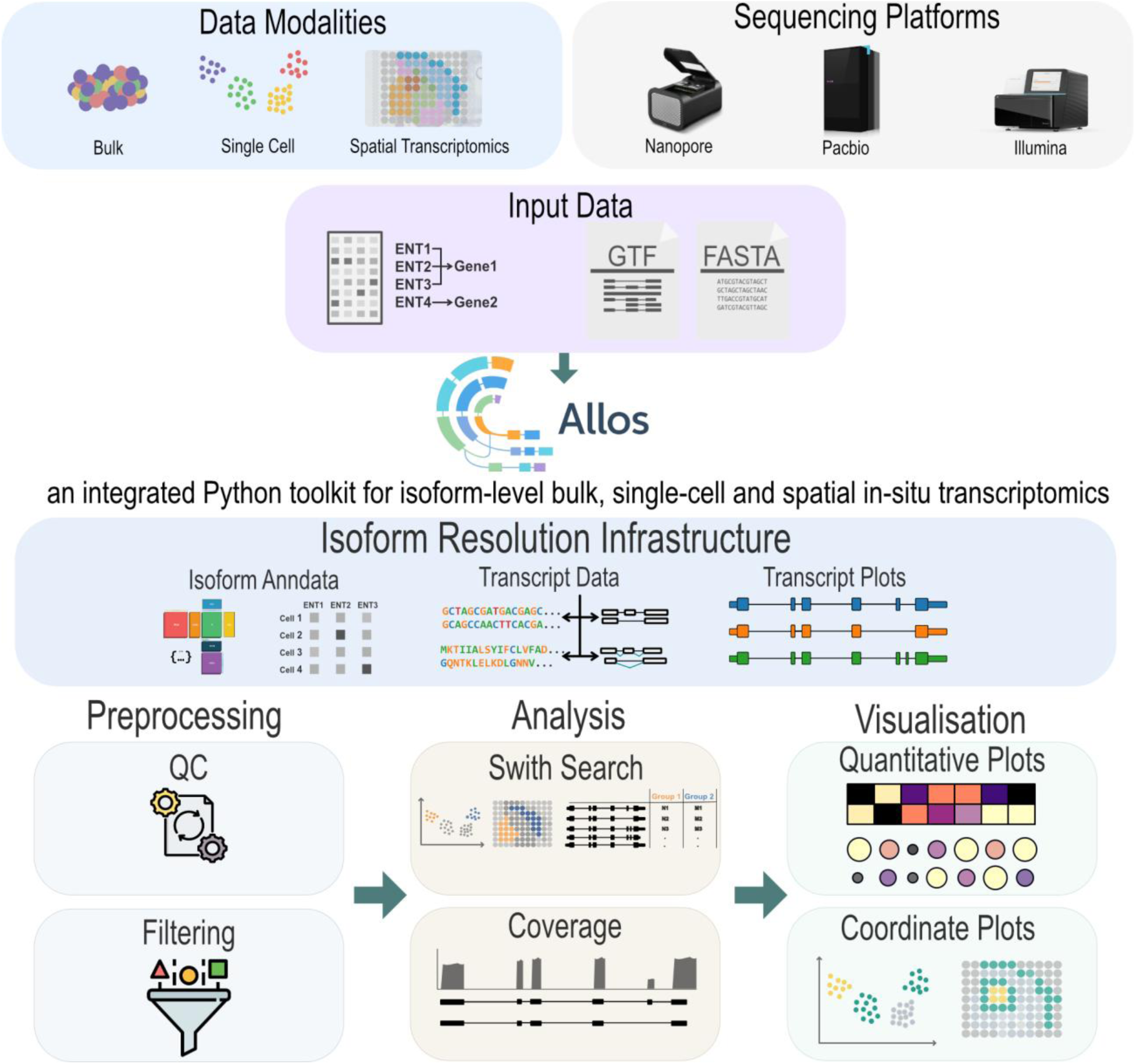
Overview of the Allos framework. Allos accepts isoform-level count matrices from bulk, single-cell, and spatial transcriptomics experiments across long- and short-read sequencing platforms, alongside GTF/GFF annotations and optional FASTA references (top). Inputs are structured into an isoform-resolution AnnData object linked to a TranscriptData annotation layer and transcript structure plots (middle). The downstream workflow (bottom) covers preprocessing; analysis (differential isoform usage screening via SwitchSearch and read-level coverage validation); and visualisation via summary plots, embeddings, and spatial maps.

**Figure 2.**
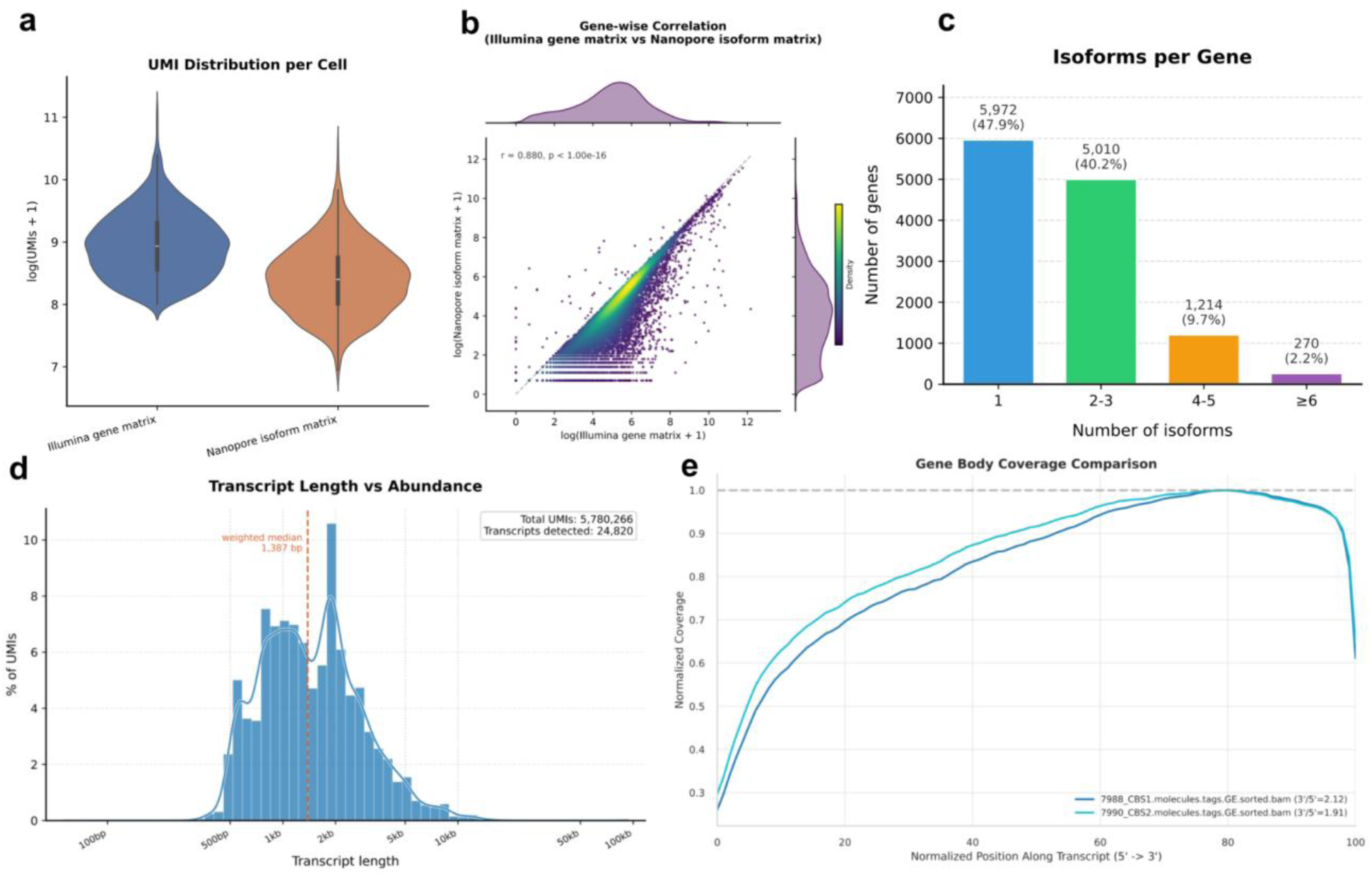
Allos QC module applied to the E18 mouse single-cell and CBS spatial datasets. (a) UMI distributions for the Illumina gene matrix and Nanopore isoform matrix. (b) Gene-wise correlation between Illumina and aggregated Nanopore isoform counts. (c) Number of detected isoforms per gene. (d) Transcript length distribution weighted by UMI abundance. (e) Gene body coverage profiles for CBS1 and CBS2 spatial replicates. Panels (a–d) are derived from the E18 dataset; panel (e) from the CBS spatial dataset.

### Demonstrating end-to-end exploration of isoform-resolved single-cell and spatial datasets

The E18 mouse brain dataset provides a useful context for demonstrating Allos’s isoform exploration capabilities. UMAP embedding (**Fig.3a**) was computed using the standard scanpy workflow on the transcript-by-cell matrix, with cells separating into expected neuronal and progenitor/glial populations. We labelled cells using the original publication labels to ensure consistency with previously reported results.

**Figure 3.**
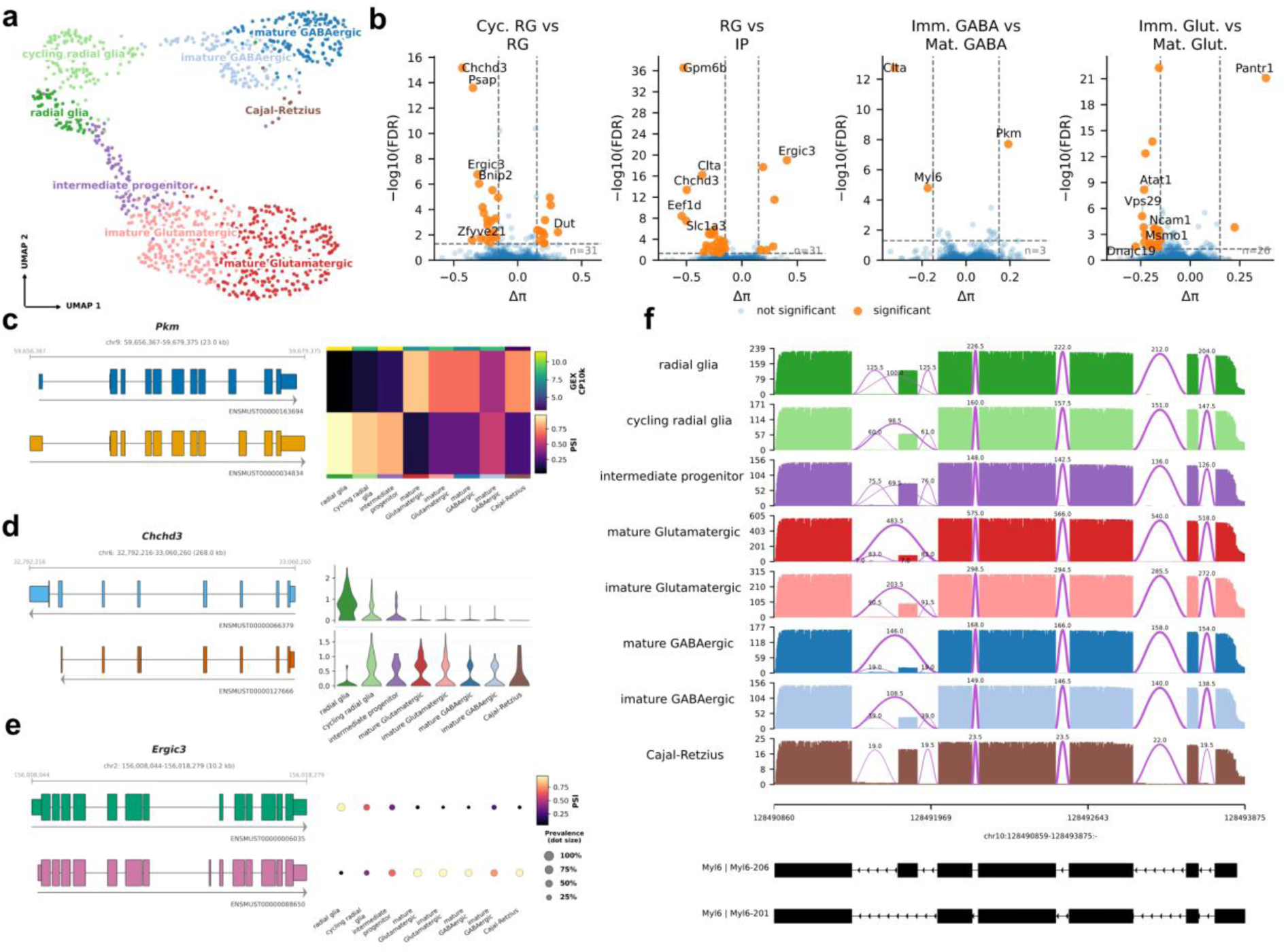
End-to-end exploration of isoform switches in single-cell data using Allos. (a) UMAP embedding generated from transcript matrix of the E18 mouse dataset, coloured by annotated cell type. (b) Representative volcano plots from pairwise SwitchSearch comparisons, showing isoform usage differences (Δπ, a summary effect size capturing the combined magnitude of the two most strongly shifting isoforms’ PSI changes) against statistical significance (−log10 FDR) for four cell-type contrasts. Highlighted genes represent significant switching events (|Δπ| ≥ 0.15, orange). (c) Composed heatmap plot for Pkm, showing transcript structures rendered to scale alongside PSI values across cell types. (d) Composed violin plot for Chchd3, showing transcript structures alongside per-cell isoform expression distributions across cell types. (e) Composed dot plot for Ergic3, showing transcript structures alongside isoform prevalence and mean PSI per cell type. (f) Coverage plot for Myl6 across cell types, showing mean read coverage and splice junction usage stratified by cell population, with reference transcript models shown below.

We apply Allos SwitchSearch across all pairwise cell-type contrasts, producing ranked volcano plots that give a transcriptome-wide view of differential isoform usage (**Fig.3b** shows a representative subset; the full set of comparisons is generated simultaneously). Significant switching events include well-characterised genes such as Pkm, Myl6, and Clta, also identified in the original publication, alongside less-studied candidates, confirming that SwitchSearch can recover known switching events and surface novel candidates across all pairwise comparisons in a single pass.

Allos supports multiple composed and standalone plot formats that jointly display transcript structure and isoform usage, allowing users to choose the representation best suited to their data. Pkm is shown using the heatmap format (**Fig.3c**). Pkm produces M1 and M2 isoforms differing by a single mutually exclusive exon, encoding distinct pyruvate kinase catalytic properties linked to oxidative versus glycolytic metabolism, and was originally reported as undergoing a pronounced cell-type-specific switch. The heatmap summarises mean PSI values across all cell types simultaneously, making it well-suited to genes with strong, consistent switching patterns. The upper expression panel confirms Pkm is broadly expressed across cell types, indicating that isoform differences reflect a true shift in splicing preference rather than differential gene expression. The PSI heatmap shows ENSMUST00000034834 (Pkm-M2) dominating in radial glia, cycling radial glia, and intermediate progenitors, while ENSMUST00000163694 (Pkm-M1) becomes dominant in mature glutamatergic and GABAergic neurons at the progenitor-to-neuron transition.

Chchd3, a core MICOS complex component required for cristae junction organisation, presents a subtler case (**Fig.3d**). Despite low overall expression, per-cell distributions reveal the exon-excluding isoform (ENSMUST00000066379) is largely restricted to radial glia, while the exon-including form (ENSMUST00000127666) is broadly expressed in neuronal populations. The alternatively spliced exon falls within the coiled-coil domain that mediates MICOS subunit interactions, suggesting the radial glia-specific isoform alters cristae organisation to meet the distinct mitochondrial requirements of neural stem cells. This pattern would be masked by pseudobulk or mean-based summaries.

Ergic3, which encodes an ER–Golgi intermediate compartment protein involved in anterograde secretory trafficking, displays a progressive developmental switch (**Fig.3e**). The exon-excluding isoform (ENSMUST00000006035) is used almost exclusively in radial glia; both isoforms co-occur in cycling progenitors, and the exon-including form (ENSMUST00000088650) becomes dominant in differentiated neurons. This graded transition parallels the expansion of the secretory pathway during neuronal maturation, as post-mitotic neurons must sustain high rates of membrane protein trafficking to build and maintain axonal and dendritic compartments. The dot plot separates the prevalence of isoform detection from its mean proportion, confirming the shift is population-wide rather than driven by a few highly expressing cells.

Coverage plots provide an orthogonal layer of read-level validation for candidate switches (71). For Myl6 (**Fig.3f**), mean read coverage and splice junction counts are shown per cell type across two technical replicates. The exon-including isoform Myl6-206 has strong junction support in radial glia, progressively decreasing through progenitor stages until Myl6-201 dominates in mature neurons. The consistency strengthens belief a genuine biological signal. Myl6 encodes the essential myosin light chain, and the alternatively included exon has been shown to modulate actin-binding affinity, suggesting the switch tunes cytoskeletal dynamics as cells transition from a proliferative, migratory state to a post-migratory morphology.

To demonstrate Allos on spatial isoform transcriptomics, we re-analysed the CBS1 and CBS2 datasets introduced above. The two sections show highly reproducible anatomical in situ spot clustering across 12 annotated brain regions (**Fig.4a**), providing a structured tissue context for assessing spatial isoform usage.

**Figure 4.**
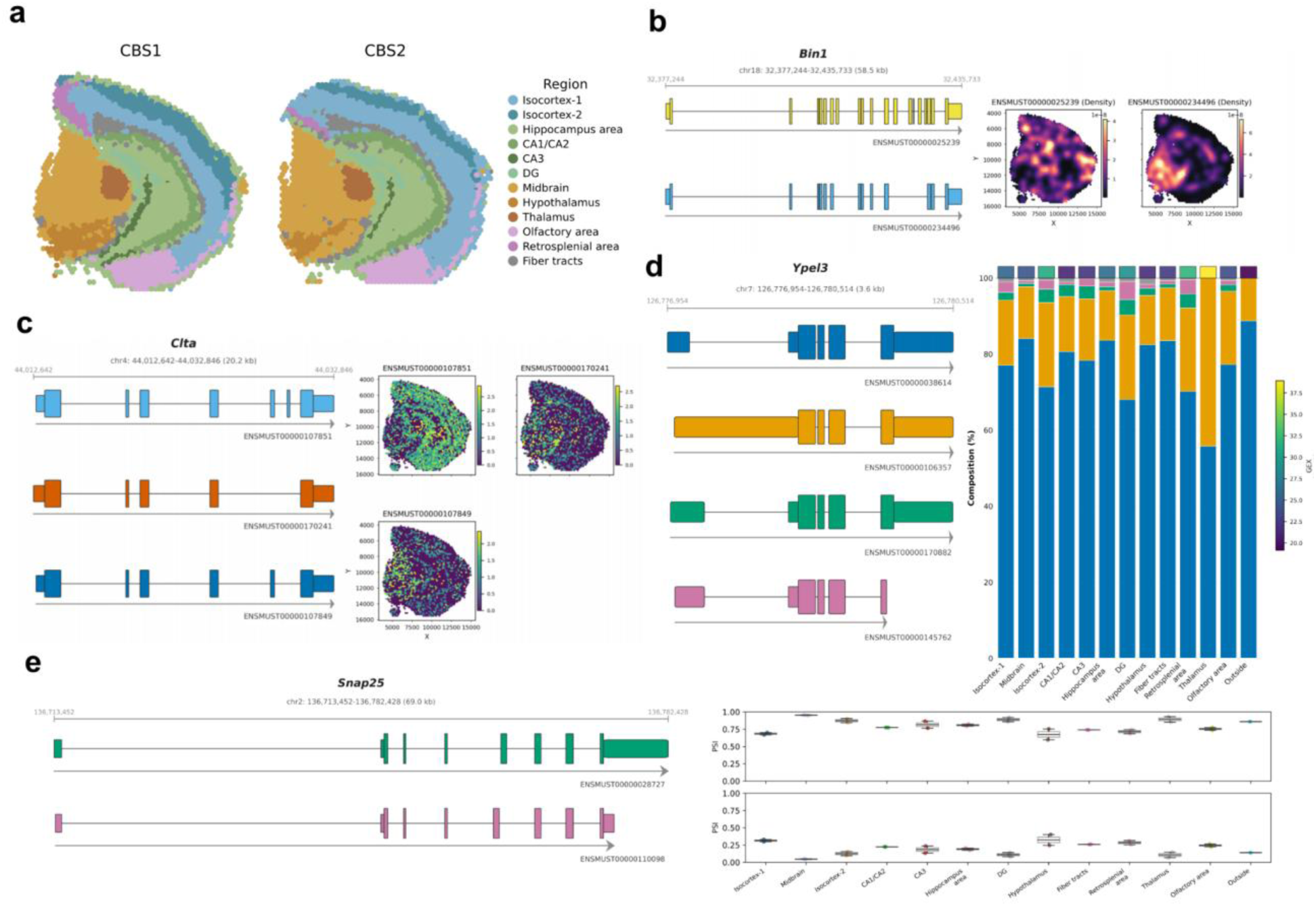
Isoform-resolution spatial transcriptomics analysis using Allos. (a) Spatial embedding of CBS1 and CBS2 coloured by annotated anatomical region. (b) Composed KDE-smoothed density plot for Bin1, showing transcript structures alongside isoform density maps across CBS1, illustrating spatially distinct enrichment patterns for two isoforms with complementary regional distributions. (c) Spot-level spatial isoform expression maps for three Clta transcripts across CBS1, showing transcript structures alongside per-spot expression values, demonstrating how simultaneous visualisation of multiple isoforms reveals distinct and complementary spatial enrichment patterns across the tissue section. (d) Composed stacked bar plot for Ypel3, showing four transcript structures alongside isoform composition across annotated regions, with colour intensity reflecting mean expression per region. (e) Replicate-resolved composed PSI plot for Snap25, showing isoform PSI distributions across all annotated regions for CBS1 and CBS2, demonstrating reproducible region-specific isoform usage across biological replicates.

We examined several genes with region-specific splicing across the CBS1 section. Bin1, a BAR-domain adaptor protein and leading Alzheimer’s disease genetic risk modifier (72), shows striking spatial isoform segregation (**Fig.4b**). ENSMUST00000025239 is broadly distributed, whereas ENSMUST00000234496 is confined to white matter and fiber tracts, consistent with the oligodendrocyte-specific expression reported in the original SiT publication. This segregation likely reflects the distinct roles of Bin1 isoforms in neuronal synaptic vesicle endocytosis versus oligodendrocyte membrane remodelling during myelination; neuronal isoforms uniquely contain a CLAP domain that mediates clathrin-dependent endocytosis, whereas the ubiquitous isoforms expressed in oligodendrocytes lack this domain(73). This highlights the importance of isoform-level resolution for interpreting cell-type-specific Alzheimer’s disease risk variants. KDE smoothing was applied to mitigate per-spot count sparsity.

Clta, encoding clathrin light chain A involved in vesicle-mediated intracellular transport, shows complementary spatial enrichment of its three detected isoforms (**Fig.4c**). ENSMUST00000107851 is broadly expressed, while ENSMUST00000170241 is enriched in distinct regional compartments. The alternatively spliced segment modulates clathrin cage assembly kinetics and Hip protein interactions, suggesting regional isoform preferences adapt endocytic dynamics to the trafficking demands of distinct neuronal populations.

Ypel3, a conserved gene implicated in glial development through zebrafish loss-of-function studies (74), shows a region-specific departure from its default isoform programme (**Fig.4d**). While ENSMUST00000038614 dominates in most brain regions, the thalamus shows both elevated Ypel3 expression and a marked increase in the relative contribution of ENSMUST00000106357. The co-occurrence of transcriptional upregulation and isoform rebalancing suggests coordinated regulation, potentially reflecting the specialised requirements of thalamic relay neurons. This pattern would be invisible to gene-level analyses, which capture only the expression increase.

Replicate-resolved PSI distributions for Snap25 across CBS1 and CBS2 are shown for all 12 annotated regions, with points representing per-replicate PSI values (**Fig.4e**). PSI values are highly consistent across replicates, supporting the conclusion that observed spatial isoform patterns reflect biological organisation rather than technical variation. This consistency is particularly notable for Snap25, whose well-characterised alternative splicing produces the Snap25a and Snap25b isoforms via mutually exclusive exon 5 usage, with established developmental and functional roles in neuronal systems (75). In multi-donor datasets, replicate divergence may additionally reflect genetic effects at splicing quantitative trait loci (sQTLs); isoform-level quantification across cell types and tissue regions in single-cell and spatial assays could therefore enable mapping of context-specific sQTLs beyond the population-level catalogues established in bulk-tissue resources such as GTEx.

### Allos enables interactive dashboards for real-time isoform exploration

Although Allos is designed primarily as a Python library, it also provides an interactive dashboard built on the Streamlit framework, exposing its core visualisation and analysis functionality through a graphical interface. This interface is inspired by popular single-cell data exploration tools such as CZ CELLxGENE, UCSC Cell Browser, and Loupe Browser, which have demonstrated the value of interactive, code-free environments for biological interpretation of complex datasets (76,77). Isoform-resolution data present a particularly compelling case for this kind of multi-resolution exploration: a single dataset can simultaneously encode variation at the level of individual transcripts, genes, cell types, developmental states, donors, replicates, spatial regions, and predicted protein products, and navigating these interacting layers of biological organisation benefits substantially from interactive visualisation by domain experts. The dashboard accepts an Allos pre-processed isoform Anndata at the isoform- or gene-level as input. It enables gene-level isoform exploration, panel configuration, and protein domain visualisation without requiring users to write code. This addresses a practical limitation in collaborative research settings, where the experts best positioned to interpret isoform-switching events are often wet-lab biologists or clinical collaborators who do not interact directly with the computational pipeline. Figures can be exported in PNG, PDF, or SVG format at publication resolution. The dashboard can be hosted locally following standard installation, and its architecture supports deployment on shared or remote infrastructure, facilitating dataset sharing within collaborative consortia.

### Allos SwitchSearch demonstrates concordance with established differential isoform usage methods

To contextualise the performance of SwitchSearch relative to established methods, we benchmarked it against DiffSplice, DEXSeq, and the previously published Sicelore results on the E18 dataset (**Fig.5a**). SwitchSearch completed the full pairwise screen in 34 seconds compared to 1.8 minutes for DiffSplice and 4.9 minutes for DEXSeq under equivalent computational conditions, demonstrating its utility as a rapid first-pass exploratory screen (**Fig.5b**). Notably, both DiffSplice, via the edgePython pseudobulk framework, and SPLISOSM are directly accessible within Allos, allowing users to move seamlessly from exploratory screening to statistically rigorous follow-up analysis in a single environment. Four-way concordance analysis confirmed substantial agreement across methods, with 52 genes identified by all four approaches supporting the reliability of the consensus hit set (**Fig.5c**). For spatial isoform data, three-way comparison with SPLISOSM across CBS1, CBS2, and their intersection revealed a consistent core of switching genes shared across methods (**Fig.5d**).

**Figure 5.**
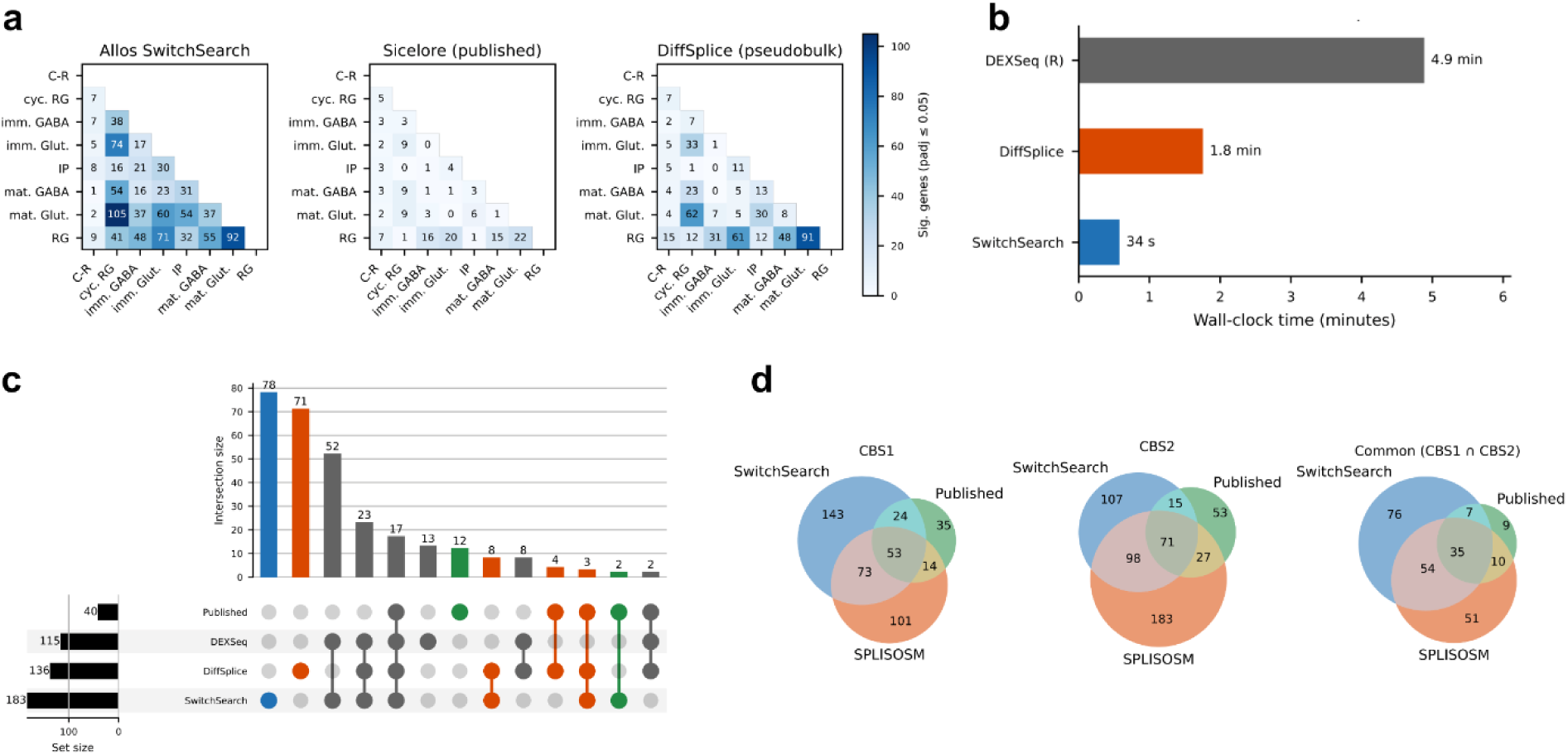
SwitchSearch benchmarking and concordance analysis. (a) Pairwise differential isoform usage across cell-type contrasts in the E18 mouse forebrain dataset for SwitchSearch, Sicelore (published), and DiffSplice (pseudobulk), displayed on a shared colour scale. (b) Wall-clock runtime comparison across SwitchSearch, DiffSplice, and DEXSeq on the E18 dataset restricted to 4 CPU cores. (c) Upset plot showing gene-level concordance between SwitchSearch (padj ≤ 0.05, |Δπ| ≥ 0.15), DiffSplice (padj ≤ 0.05, |ΔPSI| ≥ 0.15), DEXSeq (padj ≤ 0.05, |ΔPSI| ≥ 0.15), and the previously published Sicelore results (padj ≤ 0.05). (d) Three-way overlap between SwitchSearch, SPLISOSM, and published results for CBS1, CBS2, and their intersection.

As SwitchSearch and SPLISOSM operate under fundamentally different statistical frameworks, the former as a region-level pseudobulk screen, the latter using kernel-based spatial independence testing that explicitly models spatial autocorrelation and isoform compositionality, divergence between their results is expected and reflects differences in the biological patterns each approach is powered to detect. SwitchSearches χ²-based design does not account for overdispersion or spatial autocorrelation, and its false-positive rate is correspondingly elevated compared with more comprehensive methods; significant hits should therefore be treated as candidates for downstream validation. This is an explicit design tradeoff: unlike pseudobulk methods such as DiffSplice, which require biological replicates to function properly as intended, SwitchSearch operates effectively on single-sample datasets, a common scenario in single-cell and spatial transcriptomics experiments, particularly those sequenced with long reads, which makes it a practical entry point for isoform switch discovery across the broadest possible range of experimental designs.

## Discussion

Long-read and hybrid single-cell protocols now produce isoform-resolved transcriptomes at scale. Yet, analysis remains fragmented: researchers must combine separate tools for annotation, quantification, structure visualisation, and differential usage, typically outside standard single-cell workflows. Allos provides a transcript-centric alternative built on AnnData that brings isoform-level representation, exploratory screening, and structure-aware visual analytics into the scverse ecosystem. This allows isoform-resolved analyses to run alongside established clustering, annotation, embedding, and spatial workflows, while linking transcript structure, quantitative summaries, read-level evidence, and protein-level interpretation within a common Python environment.

Allos is designed as an integrated framework for representing, visualising, and prioritising isoform-level variation in single-cell and spatial transcriptomics. Its outputs necessarily inherit limitations of upstream quantification and single-cell long-read data. First, Allos is agnostic to the quantification pipeline and therefore reflects pipeline-specific choices in read assignment, transcript collapsing, barcode/UMI handling, and reference annotation; differences in these steps can change apparent isoform usage, particularly for low-abundance isoforms or highly similar transcript models. Second, annotation incompleteness and transcript-model uncertainty remain substantial challenges: sparse per-cell coverage and variable transcript completeness can lead to ambiguous isoform calls, especially when isoforms share most exons or differ primarily at transcript ends. Allos mitigates this by supporting junction- and coverage-based inspection, but these evidence layers cannot always fully resolve all ambiguities without additional validation. Finally, protein- and domain-level summaries depend on external reference resources (e.g., UniProt/InterPro/Pfam) and on CDS annotations, which can be incomplete, isoform-inconsistent, redundant, or version-dependent. Consequently, protein feature plots should be interpreted as hypothesis-generating and cross-checked against existing functional literature for the gene/protein (and, where available, supported by orthogonal evidence such as conserved domain predictions, experimental or computationally derived structural data, or proteomics validation). The structured representations that Allos maintains for each isoform (full-length transcript sequences, splice junction coordinates, CDS-derived protein translations, domain annotations, and quantitative usage) constitute the input layer needed to connect isoform discovery with biological foundation models. Tools for predicting splicing from sequence(78), inferring protein structure from translated isoform products (79), and generating functional hypotheses from protein sequence and domain architecture (80) are individually powerful, but their utility for isoform biology depends on structured, queryable access to transcript and protein data. By organising these representations into a single AnnData-compatible object, Allos provides the infrastructure to route candidate-switching events to these models at scale. In the future, we will be actively pursuing these integrations as long-read datasets grow to encompass thousands of switches across cell types, tissues, and disease states.

## Conclusion

Allos is a Python-native framework for transcript-centric representation, differential isoform usage screening, and structure-aware interpretation in single-cell and spatial transcriptomics. Built on the AnnData data model and integrated with the scverse ecosystem, it supports isoform-resolved analyses alongside established gene-level workflows within a single data format. Using long-read single-cell and spatial datasets, we demonstrate end-to-end exploration of isoform diversity from transcriptome-wide switch screening to structure-aware visualisation, coverage-based validation, and protein-level interpretation. Allos is publicly available at https://github.com/cobioda/allos.

## Methods

### Overview

Allos is implemented in Python and designed as an extensible framework for isoform-level analysis of single-cell and spatial transcriptomics data. Its modules are built around the AnnData data model and integrate transcript annotations, isoform quantification, junction-level evidence, and multiple visualisation strategies. The framework supports transcript- and gene-level analyses, differential isoform usage screening, and protein-level interpretation.

### Data representation and core object model

Allos adopts the AnnData format for seamless integration within the scverse ecosystem, providing a transcript-centric framework for analysing isoform-level expression data from long-read, plate-based, or spatial transcriptomics pipelines. It can also function as a standalone tool for structural visualisation of known and novel transcripts alongside quantitative measurements. The framework requires a transcript-level count matrix and a gene-to-transcript mapping; a GTF or GFF annotation is needed for structure visualisation, and an optional transcript FASTA enables sequence-based analyses and assessment of isoform-switching consequences. Allos supports both reference-annotated and novel transcripts. Unannotated isoforms from long-read pipelines are handled identically to known annotations across all analysis and visualisation modules, provided they appear in the input GTF. Transcript counts are stored in an AnnData object, with .var indexed by transcript ID, and gene IDs are held in a separate column. Gene-level count matrices can optionally be integrated to support joint analyses and quality control via consistency checks between gene-level expression and summed transcript abundances, enabling scalable exploration of splicing patterns within standard AnnData workflows.

### TranscriptData module

The *TranscriptData* module provides a fast, structured interface for accessing and manipulating transcript annotations and is built on top of pyranges (81), a Python library for genomic interval operations. Its design was informed by the integrated transcript annotation infrastructure developed within Isosceles, which demonstrated the value of tightly coupling transcript models with downstream single-cell quantification and analysis within a unified data model, an approach TranscriptData adapts for the Python/AnnData ecosystem (53). It parses GTF/GFF annotations into objects indexed by transcript and gene identifiers, enabling rapid retrieval of exon coordinates, strand information, CDS boundaries, and transcript-level metadata for downstream visualisation and analysis. The interval-aware pyranges backend handles tasks such as merging overlapping exons, computing genomic spans, and mapping transcript structures to a shared coordinate system. When a reference FASTA is provided, TranscriptData also supports sequence extraction for transcripts or CDS regions, linking structural annotation to nucleotide- and protein-level analyses. It forms the backbone of Allos’ transcript structure visualisation and isoform-aware analysis workflows.

### Transcript Visualisation Framework

Allos includes transcript structure visualisation tools for comparing alternative isoforms within a gene. The design draws on established representations from the R ecosystem, notably ggtranscript and IsoformSwitchAnalyzeR, adapting their visual grammar (scaled exon/intron rendering, CDS distinction, and multi-transcript overlay) into a Python-native, Matplotlib-based implementation that works directly with AnnData workflows (57,82). Transcripts are rendered to scale using genomic coordinates, allowing users to inspect exon skipping, intron retention, and alternative splice-site usage. Where CDS annotations are available, coding regions and UTRs are distinguished, with UTRs drawn at reduced height. Plots can optionally include a genomic ruler, strand arrows, and transcript labels with configurable placement. For genes with long introns or dense exon structures, introns can be compressed, and exons rescaled with minimum-width constraints to keep things legible. Row spacing, exon height, length bars, and tick marks are all adjustable. Rendering in Matplotlib keeps the stack lightweight and easy to customise. Since Matplotlib underpins much of the Scanpy/scverse plotting layer, Allos figures slot naturally into existing workflows and can be edited using standard primitives.

### Isoform-level preprocessing and quality control

Allos includes a preprocessing module for harmonising isoform-resolution AnnData objects with external single-cell datasets and performing first-pass quality control before downstream analysis. Where an isoform-level AnnData and a matched short-read gene-level dataset exist for the same experiment, Allos can align cell identities and transfer per-cell annotations, including cell type labels, batch, donor, condition, and clustering assignments, from a source to a target object. This allows curated short-read annotations to be propagated onto isoform-resolution matrices without re-running clustering or classification.

To enable comparisons with gene-level datasets, Allos implements transcript-to-gene aggregation, producing a gene matrix compatible with standard analysis pipelines. This aggregated view supports dataset-level summaries, such as detected genes per cell, and cross-platform consistency checks. A gene-level correlation QC module assesses agreement between the isoform-resolution matrix and a matched gene-level dataset, and can also be used to compare signal recovery across modalities, such as spatial versus single-cell long-read data.

### Differential isoform usage screening

Allos takes a tiered approach to differential isoform usage, from rapid exploratory screening to replicate-aware statistical testing and spatially informed inference.

SwitchSearch implements a lightweight gene-level screen using a χ² contingency framework. Cells are partitioned into user-defined groups based on covariates such as cell type, condition, or composite strata, and for each pairwise or one-vs-rest comparison, transcript counts are aggregated per group and tested using a χ² test on an n × 2 contingency table, where n is the number of expressed isoforms. Genes with fewer than two detected isoforms are excluded, and a minimum count threshold limits instability from low-support transcripts. P-values are corrected using Benjamini–Hochberg FDR. To aid ranking, Allos reports group-average PSI values, ΔPSI, and an optional Δπ effect size (76), defined as the sum of PSI changes for the two most strongly shifting isoforms, signed by the dominant direction of change.

SwitchSearch is designed for speed and accessibility, analogous to marker gene identification in Scanpy or Seurat, and shares similar limitations: its χ²-based design does not account for overdispersion or biological variability, so significant hits should be treated as candidates rather than confirmed switching events. Unlike pseudobulk methods, it operates on single-sample datasets, which is common in long-read transcriptomics, making it a practical entry point across a wide range of experimental designs.

For more rigorous inference, Allos incorporates pseudobulk-based differential splicing via edgePython (83), a Python implementation of edgeR. Aggregating transcript counts at the replicate level restores the replicate as the unit of inference, reducing cell-level sparsity and pseudoreplication. The diffSplice framework supports generalised linear modelling with empirical Bayes dispersion estimation, covariate adjustment for batch, donor, and sex, and flexible contrasts for complex study designs. Together with SwitchSearch, this covers both rapid hypothesis generation and statistically principled confirmatory analysis.

For spatial data, Allos integrates with SPLISOSM (84), which uses kernel-based spatial independence testing to detect spatially variable isoform usage while accounting for autocorrelation and compositionality. Rather than wrapping SPLISOSM directly, Allos provides a conversion utility that transforms transcript-level AnnData objects into the count tensors and coordinate tables it requires. For high-resolution platforms such as Visium HD, where per-spot counts are extremely sparse, optional spatial binning aggregates neighbouring spots into super-spots, improving count depth and inference stability while preserving tissue architecture.

### Quantitative plots for exploratory data analysis

Several plot types (heatmaps, dot plots, violin plots, and stacked bar plots), familiar from gene-level single-cell analysis, are adapted here for isoform-level data, each offering a distinct view of a switching event. Isoform proportions are computed via pseudobulk aggregation, with smoothing options to stabilise low-count estimates, and users can display either the top N isoforms by mean abundance or a custom transcript set. Heatmaps show PSI-like proportions across groups with optional annotation tracks and configurable colour scaling. Dot plots encode mean proportion as colour and cell-level prevalence as dot size, helping distinguish widespread from sparse usage, particularly useful in long-read data where per-cell counts are often low. Violin plots show the full per-cell distribution of absolute expression, with optional overlaid points for individual variability. Together, these formats address the key dimensions of an isoform switch: which isoforms are changing, in which direction, across which groups, and whether the shift is population-wide or confined to a subset of cells.

### Coordinate-based visualisation

Allos supports embedding-based visualisation of isoform expression and PSI values on low-dimensional embeddings such as UMAP, with optional density smoothing to reduce dropout-driven noise common in long-read data. Multiple isoforms can be shown as multi-panel grids with shared axes for direct comparison across the embedding. For spatial datasets, isoform usage is visualised in tissue coordinates as either spot-level scatter plots or KDE-smoothed maps, with multi-panel layouts supporting side-by-side isoform comparisons within the same section.

### Composed plots

Composed plots combine transcript structure diagrams with quantitative summaries (heatmaps, dot plots, and stacked bar plots) in a single aligned figure, directly linking structural differences between isoforms to their usage patterns across cell types, conditions, or spatial regions. The coordinate-based plots described above can also be composed in this way, embedding UMAP or spatial views alongside transcript structure for a unified view of both where and how isoforms switch. All composed plots are configurable for which panels are shown, isoform and group ordering, colour scales, and additional annotation bands.

### Coverage-based validation plots

Allos provides coverage-based visualisation of aligned reads from BAM files, aggregated across user-defined groups and displayed alongside annotated transcript structures. This genome-browser-style view lets users assess whether isoform usage and switching events are supported by the underlying data and is particularly useful for investigating genes with complex splicing, intron retention, or uncertain annotations, where discrepancies may indicate missing exons, unannotated junctions, or artefactual transcript models.

### Protein-level visualisation (protein plots)

When CDS annotations and a reference FASTA are available, Allos translates transcript sequences into predicted protein products for comparison of length, composition, and domain structure across isoforms. Protein plots highlight gains or losses of domains, truncations, and frame-altering events from alternative splicing. Domain annotations can be retrieved from Ensembl and UniProt and overlaid onto transcript diagrams, directly mapping exon-level differences to predicted protein consequences (85,86).

## Declarations

### Ethics approval and consent to participate

Not applicable.

### Consent for publication

Not applicable.

### Availability of data and materials

All datasets analysed in this study are publicly available. The E18 mouse single-cell dataset is available from Gene Expression Omnibus (GEO) under accession number GSE130708. The mouse coronal brain section spatial isoform transcriptomics datasets (CBS1 and CBS2) are available from GEO under accession number GSE153859. Allos is open-source software released under the MIT licence and is freely available at https://github.com/cobioda/allos. The package can be installed directly from the repository using: pip install git+https://github.com/cobioda/allos.git. Comprehensive documentation, tutorials, and example notebooks are provided in the repository.

## Author’s contributions

K.L. supervised the project. P.B., G.V. and K.L. secured funding. E.M. and A.D. wrote the code. E.M. wrote the manuscript with input from all authors. All authors approved the final paper.

## Use of AI and AI-assisted technologies

ChatGPT (OpenAI, USA) and Claude (Anthropic, USA) were used to improve readability and language, and to assist in refining code, particularly refactoring. After using these tools, the lead author reviewed and edited all content as needed and takes full responsibility for the publication’s content.

